# Differences in resources use lead to coexistence of seed-transmitted microbial populations

**DOI:** 10.1101/560367

**Authors:** G Torres-Cortés, BJ Garcia, S Compant, S Rezki, P Jones, A Préveaux, M Briand, A Roulet, O Bouchez, D Jacobson, M Barret

## Abstract

Seeds are involved in the vertical transmission of microorganisms in plants and act as reservoirs for the plant microbiome. They could serve as carriers of pathogens, making the study of microbial interactions on seeds important in the emergence of plant diseases. We studied the influence of biological disturbances caused by seed transmission of two phytopathogenic agents, *Alternaria brassicicola* Abra43 (Abra43) and *Xanthomonas campestris* pv. *campestris* 8004 (Xcc8004), on the structure and function of radish seed microbial assemblages, as well as the nutritional overlap between Xcc8004 and the seed microbiome, to find seed microbial residents capable of outcompeting this pathogen. According to taxonomic and functional inference performed on metagenomics reads, no shift in structure and function of the seed microbiome was observed following Abra43 and Xcc8004 transmission. This lack of impact derives from a limited overlap in nutritional resources between Xcc8004 and the major bacterial populations of radish seeds. However, two native seed-associated bacterial strains belonging to *Stenotrophomonas rhizophila* displayed a high overlap with Xcc8004 regarding the use of resources; they might therefore limit its transmission. The strategy we used may serve as a foundation for the selection of seed indigenous bacterial strains that could limit seed transmission of pathogens.

## INTRODUCTION

Plant-associated microbial assemblages (a.k.a plant microbiome) can impact plant fitness through the modification of a number of traits including biomass production, flowering time or resistance to abiotic and biotic stresses^1–4^. Due to these important effects on plant growth and health, numerous studies have focused mainly on the microbial communities horizontally acquired from the environment and associated with the rhizosphere and phyllosphere^5–7^. Although the relative contribution of vertical transmission in the ultimate composition of the plant microbiome is probably limited^8^, vertically-transmitted microorganisms can also have an important impact on plant fitness^9^. For instance, vertically-transmitted microorganisms can promote germination of seeds through the production of cytokinins^10, 11^ or increase nutrient availability in shoots and roots^12^. A well-documented example of vertical transmission is seed transmission of plant pathogenic agents. Seed transmission of plant pathogens, including viruses, bacteria and fungi, was initially identified in the late 1800s - early 1900s^13^. Seed-transmission of plant pathogenic agents serves as a major means of dispersal and can therefore be responsible for disease emergence and spread^13, 14^. Very low degrees of seed contamination by bacterial pathogens can lead to efficient plant colonization^15^. Furthermore, seed transmission of plant pathogens is usually asymptomatic and can take place on non-host plants^16, 17^, which can then serve as a reservoir of plant pathogens.

To control seed transmission of plant pathogens one should either eradicate them on seed-producing crops or perform seed treatments. Fungicide application during plant production or as seed treatment is an effective means of fungal pathogen management^18^. Since these chemical-base methods are unsatisfactory for bacterial plant pathogens and have a potentially harmful environmental impact^19^, alternative control methods have been proposed, including seed health testing, physical seed treatment (e.g thermotherapy) and biological methods such as seed coating of specific biocontrol agents^14, 18^. Direct incorporation of biocontrol agents within the seed tissues through inoculation of seed-producing crops is of interest to restrict seed transmission of plant pathogenic agents. Recently, this approach (called EndoSeed^TM^) has been successfully performed by introducing the plant-growth promoting strain *Paraburkholderia phytofirmans* PsJN into seeds of multiple plant species through spray-inoculation of the parent plant at the flowering stage^20^. Therefore, inoculation of strain(s) that can compete with plant pathogens for resources and space may be of interest to limit seed transmission of phytopathogenic agents^21^.

Although inoculation of biocontrol microorganisms on or within the seeds may improve seedling health by protecting them against seed-borne or soil-borne pathogens, adequate formulation of these microorganisms for a successful seed colonization and subsequent persistence on seedlings is still challenging^18^. Further knowledge about the interactions between phytopathogenic agents and other members of the plant microbiome could provide clues on the selection of candidate strains that compete with the target plant pathogen. These microbial interactions can be estimated by analyzing the response of the seed microbiome following seed-transmission of plant pathogens. Indeed, this approach can potentially reveal co-occurence and more importantly exclusion between plant pathogens and resident members of the seed microbiota^22^. Recently, we initiated such an approach by studying the response of the radish seed microbiome to seed-transmission of two plant pathogens of brassicas, the bacterial strain *Xanthomonas campestris* pv. *campestris* 8004 (Xcc8004^23^) and the fungal strain *Alternaria brassicicola* Abra43 (Abra43^24, 25^). These plant pathogens are frequent seed colonizers of a range of Brassicaceae species including cabbage, cauliflower, turnip and radish^26, 27^. Additionally, they differ in their seed transmission pathways: Xcc8004 being seed-transmitted through the systemic and floral pathways while Abra43 being transmitted through the external pathway^28–30^.

According to community profiling approaches, seed transmission of Abra43 impacted the taxonomic composition of the fungal fraction of the seed microbiome, while Xcc8004 did not change the structure of the seed microbiota^24^. The absence of changes in community structure following Xcc8004 seed transmission could be explained by differences in ecological niches between the plant pathogen and the members of the seed microbiome. In this work, we therefore investigated competition for resources (nutrient and space) between Xcc8004 and seed-borne bacterial strains (epiphytes and endophytes) of radish (*Raphanus sativus* var. Flamboyant5) through a combination of metagenomics, metabolic fingerprinting, bacterial confrontation assays, and fluorescence *in situ* hybridization approaches.

## RESULTS

### Impact of Xcc8004 and Abra43 transmission on the structure of the seed microbiome

The relative abundances (RA) of Xcc8004 and Abra43 within the radish seed microbiome were assessed through mapping of the metagenomic reads against their reference genomic sequences. According to mapping results, Xcc8004 RA increased from 1% of paired-end reads in the control samples to 6% and 30% for X2013 and X2014, respectively (**Additional file 1)**. Abra43 RA increased from 0.1% to 1.1% in 2013 and from 0.02% to 0.58% in 2014 (**Additional file 1)**.

Metagenomic reads were used to predict the structure of the seed microbiome with a modified and parallelized version of Kraken (ParaKraken). The vast majority of the classified reads were derived from two bacterial families: *Erwiniaceae* and *Pseudomonadaceae* (**Fig. 1a**), while less than 0.06% of reads were classified as *Dikarya* (**Additional file 2**). At a genus level (**Fig. 1g**), the seed microbiome was mainly composed of *Pantoea* and *Pseudomonas*. In addition, we detected an increase in the RA of reads affiliated with *Xanthomonas* and to a lesser extent *Alternaria* in seed samples contaminated with Xcc8004 and Abra43, respectively (**Fig. 1a** and **Fig. 1g**).

**Figure 1:**
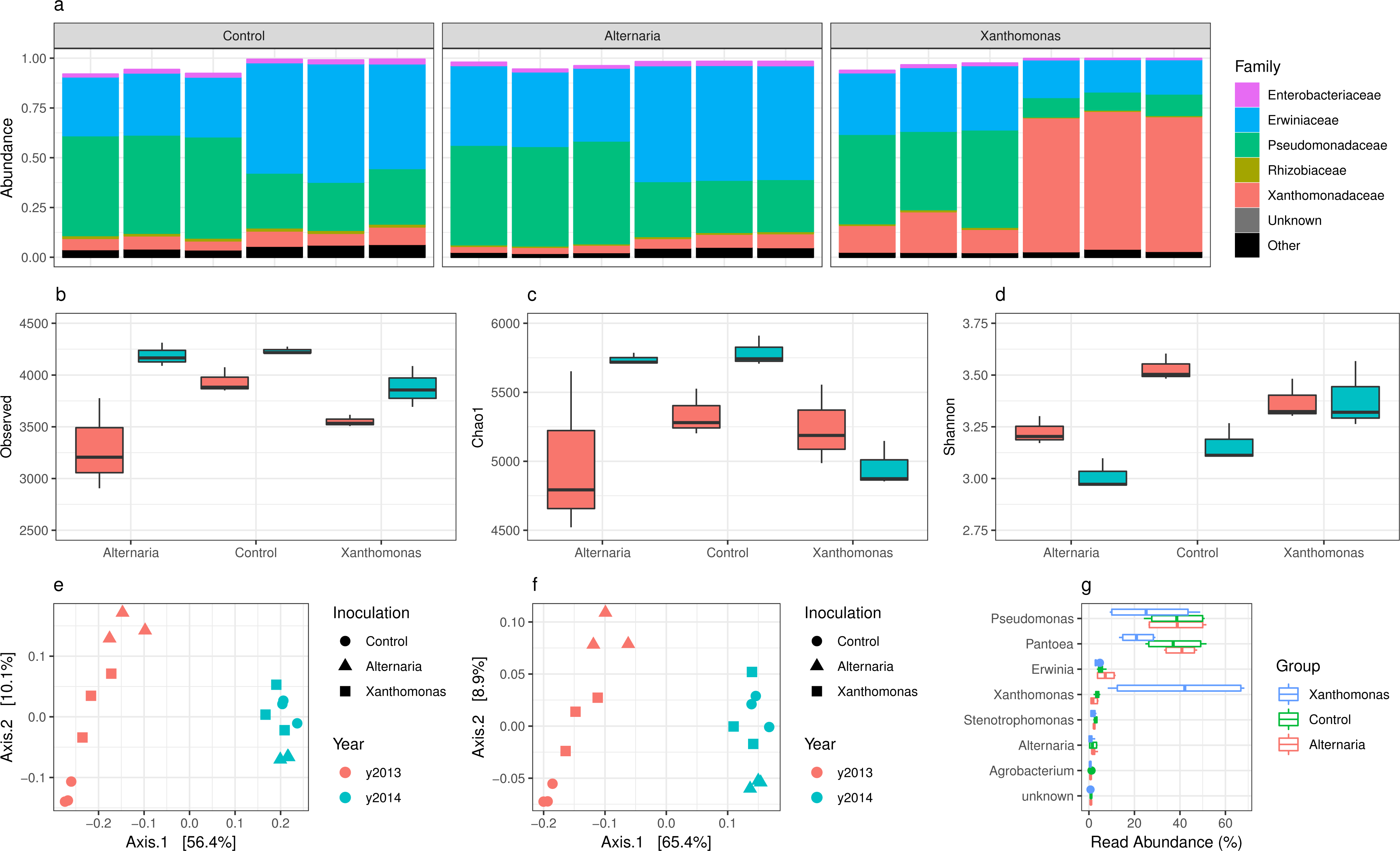
Changes in seed microbiome structure following seed transmission of Abra43 and Xcc8004. ParaKraken prediction of the taxonomic composition of the bacterial fraction of the radish seed microbiome **(a).** Measure of microbial alpha diversity through observed richness (number of predicted species), estimated richness (Chao1 index) and diversity (Shannon index) **(b-d)**. Similarity in community membership (Jaccard index) **(e)** and community composition (Bray-Curtis index) **(f)** between seed samples. Changes in relative abundance of the 10 most abundant microbial genera following seed transmission of phytopathogenic agents **(g)**.

To assess the impact of Xcc8004 and Abra43 transmission on the seed microbiome structure, we first removed the reads affiliated with the species *Xanthomonas campestris* and *Alternaria brassicicola* in the species table generated with ParaKraken. On average, 3,850 species (SD ± 385) were detected in seed-associated microbial assemblages collected from control plots (**Fig. 1b**). This estimation is one order of magnitude higher than the richness predicted with 16S rRNA gene or *gyrB*^24^. According to Kruskal-Wallis non parametric analysis of variance followed by Dunn’s post-hoc test, the observed species richness, the predicted species richness and the species diversity were not significantly affected (*P* > 0.01) by seed transmission of both phytopathogenic agents (**Fig. 1b**). Differences in community membership and community composition between seed samples were assessed with Jaccard and Bray-Curtis indexes, respectively (**Fig. 1e** and **Fig.1f**). While seed transmission of Xcc8004 and Abra43 did not alter community membership and composition (*P* > 0.01), the harvesting year explained 54.9% (*P* = 0.001) and 62.8% (*P* = 0.001) of variation in community membership and composition, respectively.

Although the overall structure of the radish seed microbiome was not impacted by Xcc8004 and Abra43 seed transmission, we assessed changes in specific taxa RA through fcros^31^ and DUO^32, 33^ analyses. High abundance species (> 0.1% RA) across all three conditions (i.e., Xcc8004 seed transmission (X), Abra43 seed transmission (A) and control (C)) were compared to identify differentially abundant species (**Additional file 3**). Of the 35 differentially abundant species across all conditions, 22 (63%) were *Pseudomonas*. DUO of the non-pathogen reads identified two networks with positive associations and no network with negative association (**Additional file 4**). The two networks were largely split by genus with one network having 67% of interacting species as *Pantoea* and the other with 67% of *Pseudomonas*. Both genera were predominant in the seed microbiome. None of the abundant species had a DUO score of >0.75, also suggesting that these pathogens had no major impact on the structure of the microbiome.

### Functional composition of the seed microbiome

Taxonomic profiling of the seed microbiomes has shown no significant differences in the community structure between samples harvested from control plots and plots inoculated with Xcc8004 and Abra43. As genomes from different isolates of the same bacterial species can display considerable genetic variation^34^, seed-transmission of these phytopathogenic microorganisms could still influence the functional composition. We therefore evaluated the impact of seed transmission of plant pathogens on the functional profile of the seed microbiome.

A total of 682,374 coding sequences (CDSs) shared in 17,589 orthologous groups (OGs) were predicted across the different metagenomic samples. On average, each metagenomic sample was composed of 8,828 OGs (SD ± 494, **Fig. 2a**). Seed transmission of Abra43 and Xcc8004 did not significantly alter (*P* > 0.01) the observed functional richness, the estimated functional richness and the functional diversity of the seed microbiome (**Fig. 2a-c**). While harvesting year significantly impacted functional membership (*P* < 0.0001, 44.4%) and functional composition (*P* < 0.0001, 52.6%) of the seed microbiome, no significant differences (*P* > 0.001) in functional membership or composition (**Fig. 2d** and **Fig. 2e**) were observed after seed transmission of Abra43 and Xcc8004. In addition, we did not detect any significant difference in RA of broad functional categories following pathogen transmission (**Fig. 2f**). Altogether these results highlight that seed transmission of Xcc8004 and Abra43 do not modify either the predicted structure or function of the seed microbiome.

**Figure 2:**
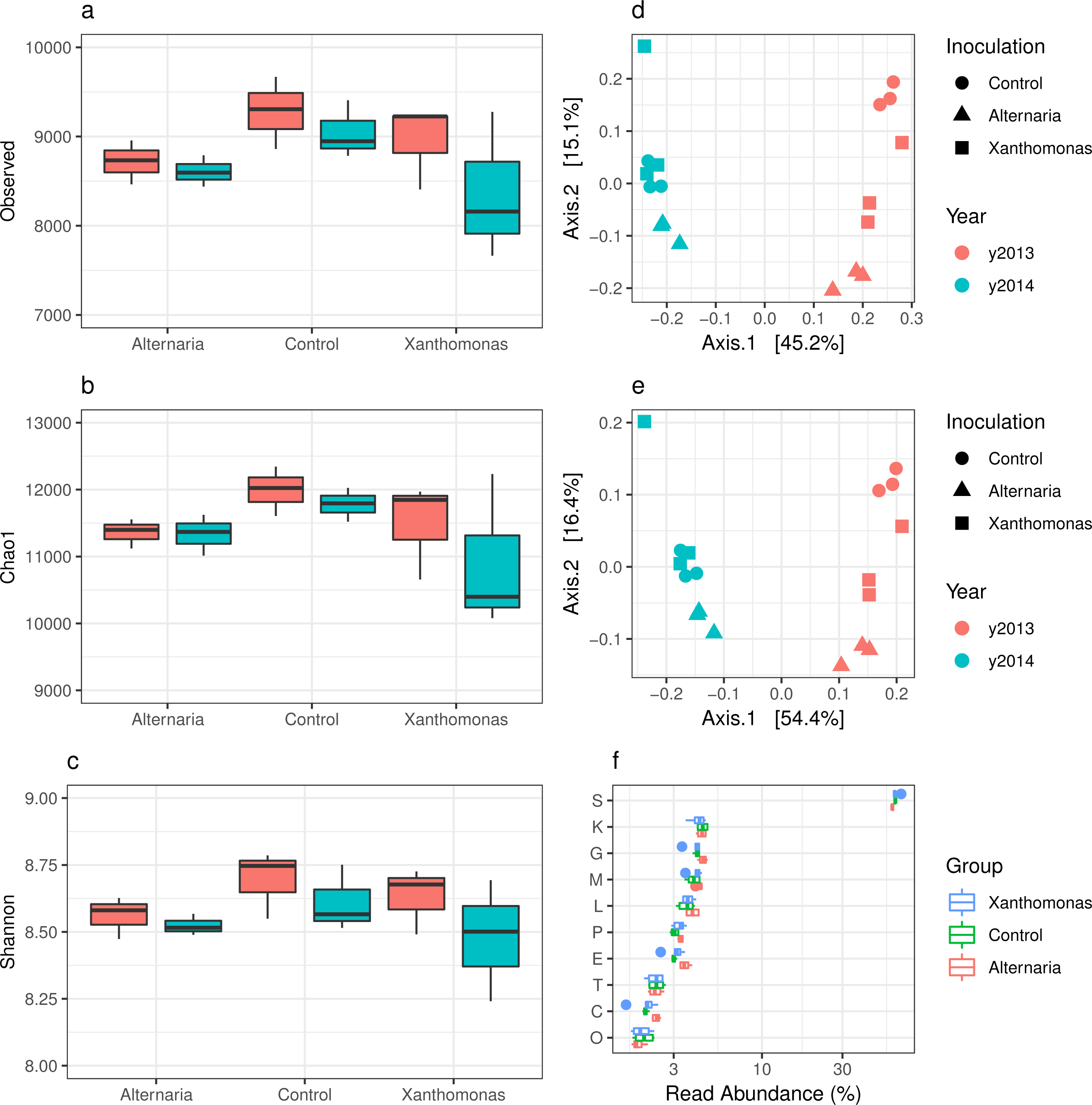
Changes in the functional composition of the seed microbiome following seed transmission of Abra43 and Xcc8004. Measure of observed functional richness (number of OG), estimated functional richness (Chao1 index) and functional diversity (Shannon index) **(a-c)**. Similarity in functional membership **(d)** and composition **(e)** as assessed with Jaccard and Bray-Curtis indexes. Changes in relative abundance of broad functional categories **(f)**.

While broad functional changes were not observed with seed transmission, OG RAs clustered together based upon relative shifts within the microbiome. Markov cluster algorithm of DUO networks resulted in 16 clusters (>2 OGs) with positive associations and 6 clusters with negative associations (**Additional file 5**). One such positive cluster is suggestive of invasion/pathogenesis of eukaryotic cells and eukaryotic cell response (**Additional file 5**), denoting important host and pathogen domains in response to infection.

### Visualization of Xanthomonas spp. within seed tissues

The absence of shift in structure and function of the seed microbiome following the plant-pathogen transmission suggests that the targeted plant pathogens and the other members of the seed microbiome do not compete for nutrient and space. We first investigated the spatial localization of Xcc8004 within seeds using fluorescence *in situ* hybridization. Using surface sterilized seeds, *Xanthomonas* spp. were detected inside inoculated seeds (**Fig. 3**). They were particularly present in embryonic tissues, especially at the surface of the radicle among other bacteria (**Fig. 3a**). They were also detected, however, in other embryonic zones of the seeds. *Xanthomonas* spp. were identified in the same tissues in control samples, but with lower abundance than Xcc8004-seeds (**Fig. 3b**). Using a photon detection method or analog integration on confocal microscope, similar results were obtained but with brighter signals from the specific probe using analog integration (**Fig. 3c** and **Fig. 3d**). Interestingly, cells were single or as packs of several bacteria (**Fig. 3a-d).** In control and Xcc8004-inoculated seeds hybridized with negative probe, no signal was identified (**Fig. 3e** and **Fig. 3f**). On the internal side of the seed coat, *Xanthomonas* spp. were further identified in Xcc8004 inoculated seeds, showing different niches of colonization (**Fig. 3g and Fig. 3h**).

**Figure 3:**
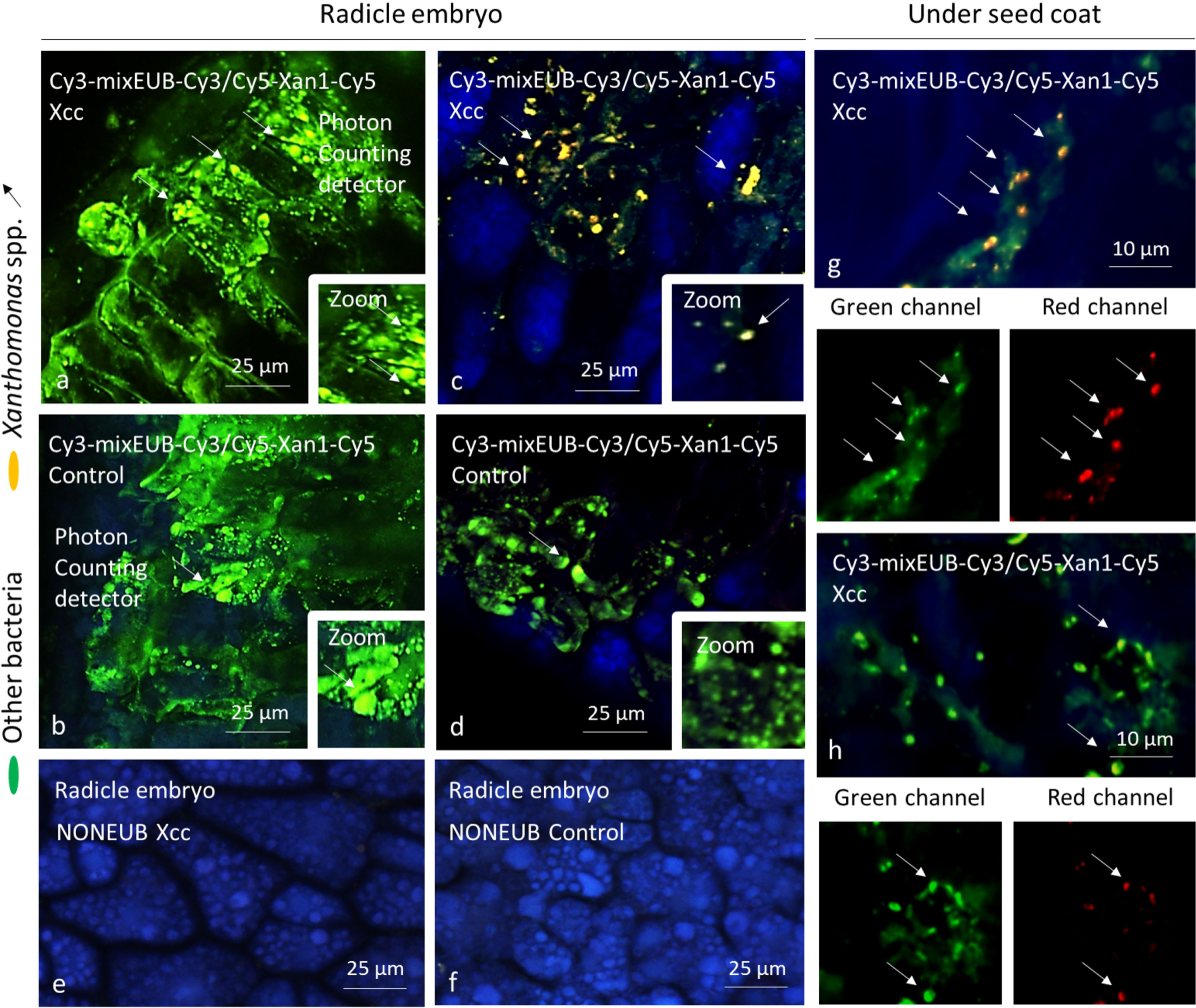
Visualization of *Xanthomonas* spp. inside seeds. Photon counting detection of *Xanthomonas* spp. on the radicle of the embryo following seed transmission of Xcc8004 (**a**) or control seeds (**b**). Analog integration detection of *Xanthomonas* spp. on radicle of seeds contaminated with Xcc8004 (**c**) or not contaminated (**d**). Use of NONEUB probes on samples subjected to Xcc8004 (**e**) or control seeds (**f**). Detection of *Xanthomonas* spp. under the seed coat (**g-h**). Bacteria corresponding to *Xanthomonas* spp. appear as yellow/orange fluorescents after blue, green and red channels were merged or as red if channels are separated while other bacteria appear as green fluorescents.

### Reconstruction of genomic sequences from the main bacterial population of the seed microbiome

Competition for nutritional resources between phytopathogenic agents and other members of the seed microbiome was subsequently estimated with genomic sequences. Given the limited number of fungal reads in our metagenomics datasets, we restricted this comparative genomic analysis to bacterial populations and Xcc8004. Bacterial genomic sequences were either recovered through metagenomic read sets or *via* genome sequencing of representative bacterial isolates (see materials and methods). Overall, 11 metagenome-assembled genomes (MAGs; >50% completeness, <10% contamination) and 21 genomes sequences were obtained (**Table 1**). These genomic sequences represented the major bacterial species detected with the ParaKraken taxonomic classification approach, and, based on their *gyrB* sequences, represented the main *gyrB* amplicon sequences variants (ASVs) of the radish seed microbiomes (**Table 1** and **Additional file 6**^24, 35^). According to the average nucleotide identity based on blast (ANIb) values, these 32 genomic sequences were divided into 22 bacterial species (**Additional file 7**). The relative abundance of each genome sequence within the seed microbiome was estimated by mapping the metagenomics reads to these sequences and expressed as average coverage. Overall, genomic sequences related to *Pantoea agglomerans* and *Pseudomonas viridiflava* were highly abundant in all radish seed samples, with an average coverage of approximately 200X and 80X, respectively (**Table 1** and **Additional file 8**).

**Table 1:**
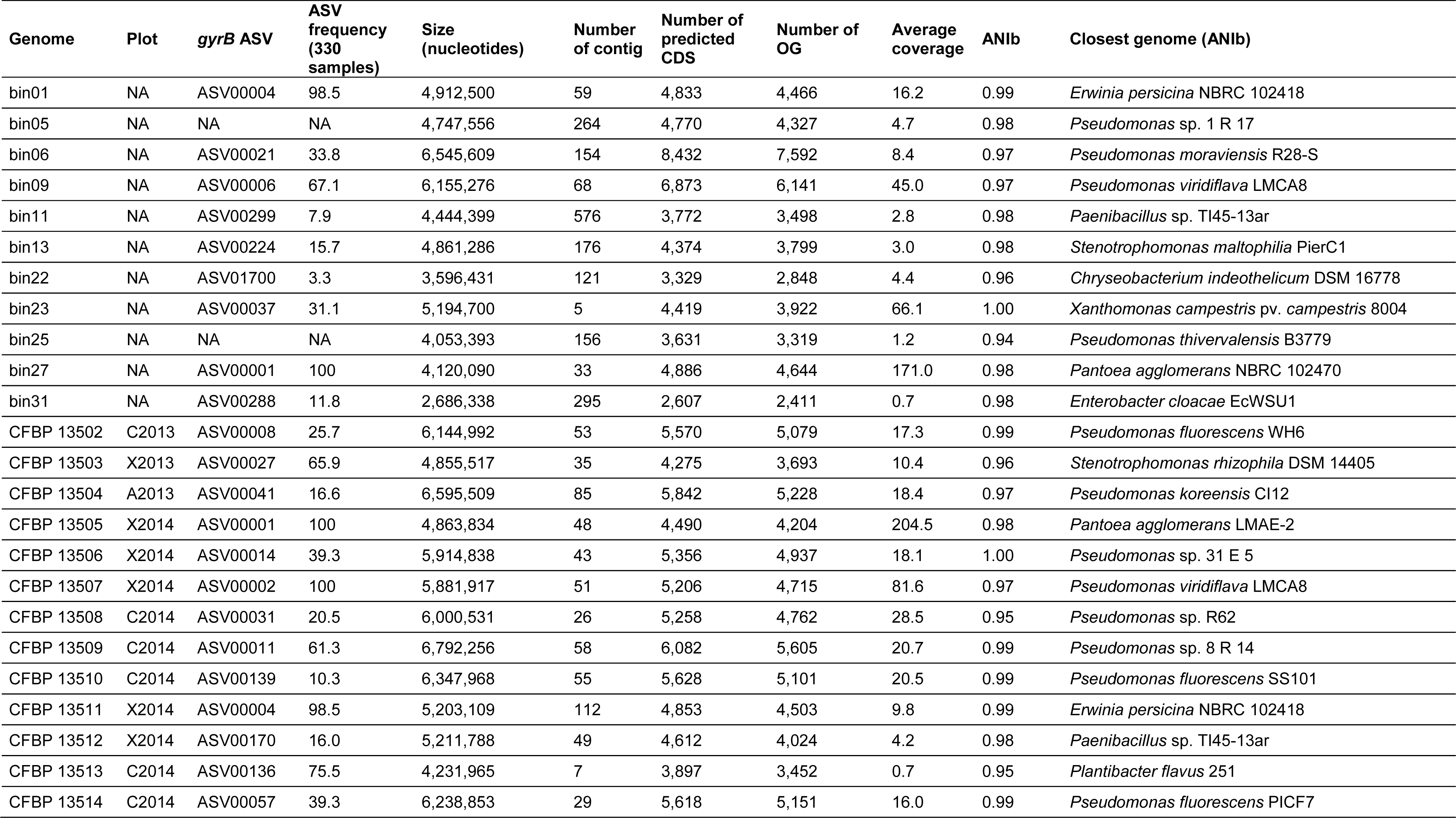
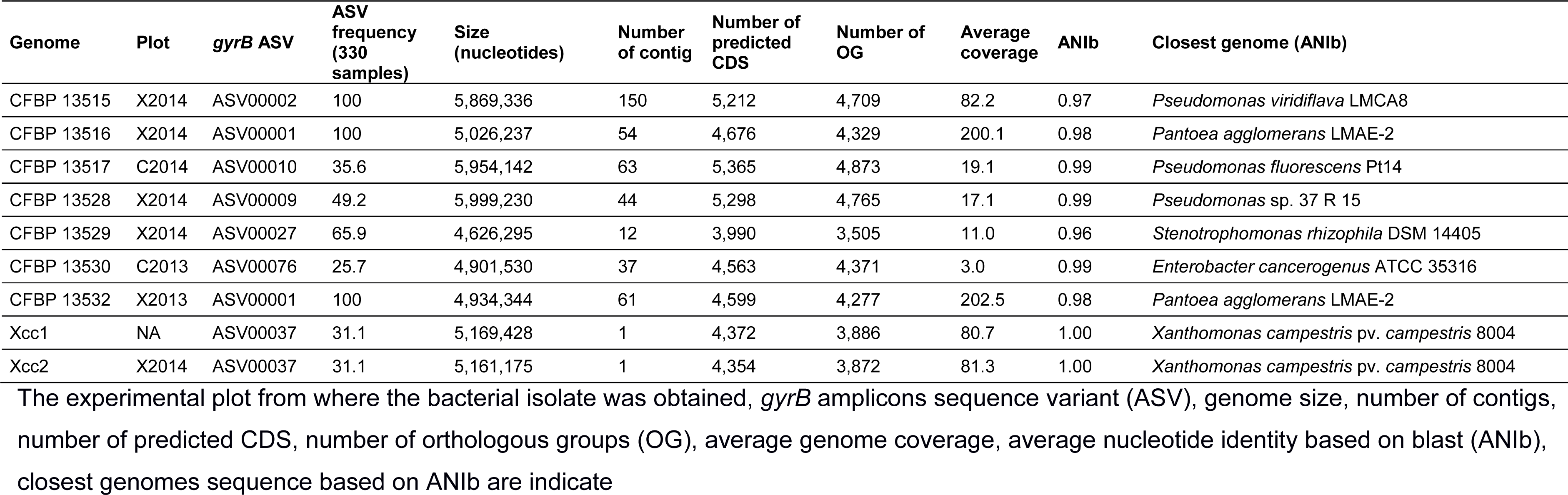
Characteristics of the genomes sequences reconstructed from metagenomic reads or sequencing of some bacterial isolates.

### Assessment of resources overlap among bacterial members of the seed microbiome

Overlap in nutritional resource use between Xcc8004 and bacterial populations from the seed microbiome was assessed by profiling nutrient consumption patterns of Xcc8004 and seed bacterial isolates with GEN III MicroPlate. Overall, the bacterial consumption pattern was largely grouped by the phylogenetic relationships between strains (**Additional file 9**). Overlap in nutritional resources between Xcc8004 and seed-associated bacterial strains ranged from 0.23 to 0.50, with the highest overlap being observed with strains CFBP13503 and CFBP13529 of *Stenotrophomonas rhizophila* (**Fig. 4**).

**Figure 4:**
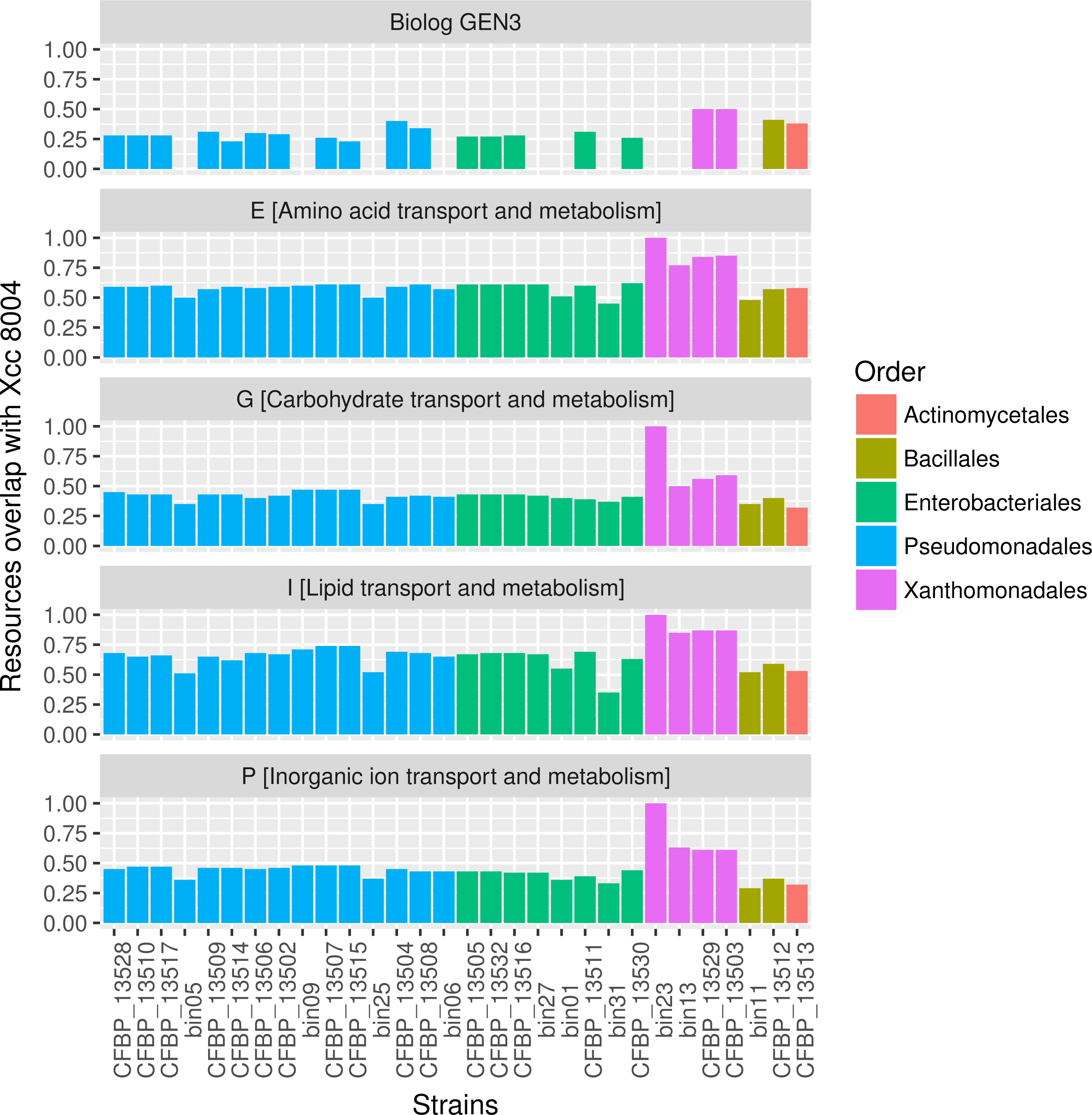
Resources overlap between Xcc8004 and bacterial populations of the seed microbiome. Resource consumption pattern was either assessed with Biolog GEN III MicroPlate or with orthologous groups (OGs) predicted from bin/genomic sequences. Resources overlap was defined as the number of shared resources between Xcc8004 and the tested bacterial strain, divided by the total number of resources used by Xcc8004 and the tested strain.

Since the compounds reduced in GEN III MicroPlates by the bacterial strains are neither representative of all seed exudates compounds nor a reflection of their actual concentrations^36^, we further assessed similarities/differences in nutritional resources consumption between Xcc8004 and the other bacterial members of the seed microbiome by using MAGs and draft genome sequences and genome sequences. We compared the number of predicted OGs associated with amino acid [E], carbohydrate [G], lipid [I] and inorganic ion [P] transport and metabolism (**Fig. 4**). The highest median overlap was observed with the functional categories E (0.61) and I (0.66), while the median overlap for G and P was below 0.5. In all cases, the most pronounced resource overlap was associated with genome sequences of *Xanthomonadales* (**Fig. 4**).

To determine if the observed resource overlap was associated with a decrease of Xcc8004 population during competition with these bacterial strains, co-inoculation of Xcc8004 was performed with each seed bacterial isolate on radish seed exudates media (see materials and methods). We observed a significant decrease (*P* < 0.01) in Xcc8004 CFU at 2 3, 4 and 5 dpi during co-inoculation with strains CFBP13503 and CFBP13529 (**Fig. 5**). This effect seems to be dependent on competition for nutritional resources, since no direct antagonism was observed between Xcc8004 and *S. rhizophila* strains during overlay assay (data not shown). To gain insight into potential competition for nutrients between Xcc8004 and *S. rhizophila* strains, we compared the set of OGs that were exclusively shared between these strains (**Additional file 10**). A total of 251 CDSs divided into 219 OGs were specifically shared between Xcc8004 and *S. rhizophila*. While, most of these OGs have no predicted function, nine CDSs were related to carbohydrate utilization (pectacte lyase, mannosidase and glucosidase). In addition, multiple protein-coding genes involved in rapid utilization of the limiting resource(s), such as TonB-dependent transporters (TBDTs) were also shared between the *Xanthomonadales* strains (**Additional file 10**).

**Figure 5:**
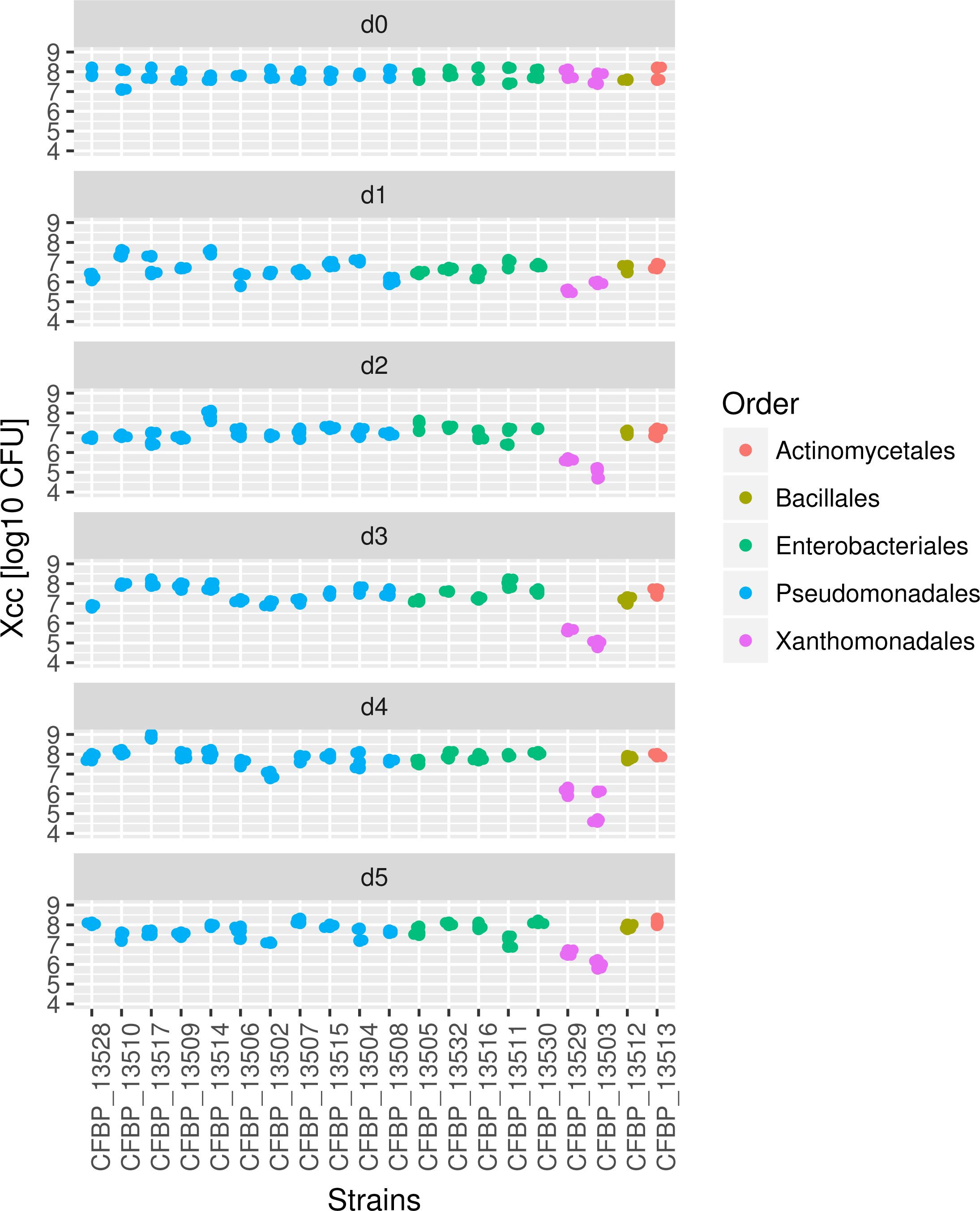
Competition for resources between Xcc8004 and bacterial populations of the seed microbiome. Competition between Xcc8004 and seed bacterial isolates was performed at a 1:1 ratio on radish exudates media. Bacterial spots were removed 1, 2, 3, 4 and 5 days post-inoculation and the number of colony forming units of Xcc8004 (*y*-axis) was recorded on TSA10% supplemented with rifampicin and X-Gluc. Three independent biological replicates, each consisting of 3 technical replicates, were performed. Differences in number of CFUs per treatment were assessed by one-way ANOVA with post-hoc Tukey’s HSD test.

### Dynamics of seed-associated microbial taxa during plant germination and emergence

Microbial populations associated with seeds could be either transient colonizers (i.e. seed-borne) or transmitted during germination and emergence to the seedling (seed-transmitted). To investigate which bacterial populations associated with radish seed samples contaminated with Xcc8004 were seed-transmitted, we performed community profiling with *gyrB* on germinating seeds and seedlings. According to the number of *gyrB* sequences detected during these early stages of the plant life’s cycle, ASVs belonging to the bacterial species *P. agglomerans*, *P. viridiflava* and species of the *P. fluorescens* group were efficiently transmitted to seedlings (**Additional file 11**). Moreover, ASVs belonging to Xcc8004 and *S. rhizophila* were also detected in germinating seeds and radish seedlings (**Additional file 11**). The competition for resources between *Xcc*8004 and *S. rhizophila* strains observed on exudates media (**Fig. 5**) and the co-occurrence of these two species within germinating seeds and seedlings either suggest that resources are not a limiting factor within these habitats or that both species are located in different tissues.

## DISCUSSION

Microbial interactions occurring on and around the seed are important for plant fitness, since seed-borne microorganisms are the primary source of inoculum for the plant. Depending on the outcome of the interactions, these microbial assemblages could either promote or reduce plant growth^9^. Understanding the processes involved in the assembly and resilience of the seed microbiota is especially relevant to comprehend the emergence of plant diseases. In the current work we assessed whether niche preemption occurred during seed-transmission of two phytopathogenic agents, Xcc8004 and Abra43, which differ in their transmission modes.

According to our metagenomic datasets, the radish seed microbiome was mostly composed of eubacteria. Whether this estimation reflects the actual structure of the radish seed microbiome or represents a bias due to DNA extraction/classification methods is currently unknown. Bacterial communities of radish seeds were dominated by two bacterial families, the *Erwiniaceae* and *Pseudomonadaceae*. More precisely, bacterial populations affiliated with *Pantoea agglomerans*, *Erwinia persicina, Pseudomonas viridiflava, P. fluorescens* and *P. koreensis* groups were associated with all radish seed samples. These species have been frequently reported in seeds of Brassicaceae^24, 35, 37–39^ and in other plant families such as Fabaceae^37^, Amaranthaceae^38^ or Poaceae^39, 40^. The high prevalence of these taxa in various seed samples may be explained by specific determinants that could help these microorganisms to thrive in the seed habitat. For instance, seed-transmitted microorganisms should deal with the high level of basal defense response related to reproductive organs^41–43^ and tolerate desiccation occurring during seed maturation^44^. In addition, the high abundance and occurrence of these bacterial populations within seed samples suggested that these taxa could influence the community structure of the plant microbiome during its early developmental stages. This assumption remained to be validated experimentally through synthetic ecology approaches as recently performed in the maize rhizosphere^45^.

In our experimental design, the impact of Xcc8004 and Abra43 transmission on the structure and function of the seed microbiome was assessed during two consecutive years. The changes in taxonomic and functional profiles were mostly driven by the harvesting year, thus confirming previous results obtained through community profiling approaches^35, 37, 46^. This is perhaps not surprising, as abiotic factors, such as soil or field management practices, have been already observed to have a strong influence in the seed microbita^37^. In contrast, the seed transmission of the two plant pathogens, Xcc8004 and Abra43, did not impact the overall composition of the seed microbiome.

Previous reports have shown that Xcc8004 is seed-transmitted through the xylem and the stigma, while Abra43 is seed-transmitted *via* fruits^30–32^. Hence Xcc8004 is probably one of the primary colonists of the seed microbiota, while Abra43 is a late-arriving species. Although both phytopathogenic agents were efficiently transmitted to radish seeds, they impacted neither the structure nor the function of the seed microbiome, implying that they are not competing with the other members of the microbiome for the same ecological niches in the seed habitat. According to *in situ* hybridization, *Xanthomonas* sp. co-occurred with other bacterial taxa within the seed coat and at the surface of the seed embryo. Since no spatial separation was observed between Xcc8004 and other seed-borne bacterial populations, then differences in resources consumption can probably explain this co-existence. Based on our genomics prediction of resource overlap and competition assays performed on radish exudate media, most of the bacterial populations of the seed microbiome are not competing with Xcc8004 for nutritional resources. Although the competition assay performed does not necessary reflect the actual composition and concentration of nutrients that are available for microbial growth during seed development^36, 49^, it can serve as a proxy for assessing competition between seed-borne bacterial species.

Under our experimental conditions, the only bacterial population that competes for nutritional resources with Xcc8004 belongs to the species *S. rhizophila*. Competition for a limiting resource can be divided into two categories: exploitative competition that is related to the rapid use of the limiting resource and contest competition that involves antagonistic interactions with production of antimicrobial compounds^50^. As we did not observe an antagonistic relationship with the overlay assay used in this work, the observed competition between *S. rhizophila* strains and Xcc8004 is likely due to resources use^48^. This contest competition is frequently related to efficient uptake of nutrients by the competing species^50^. Interestingly numerous OGs that are specifically shared between Xcc8004 and *S. rhizophila* strains CFBP13503 and CFBP13529 are related to TBDTs, which form a specific carbohydrate utilization system^51^.

According to the community profiling approach performed on more than 300 radish seed samples^24, 35^, it appears that *S. rhizophila* strains and Xcc8004 co-exist within the seed habitat. Moreover, both bacterial species co-occurred on germinating seeds and seedlings during germination and emergence, suggesting that nutritional resources are not limited enough within these habitats for observing a strong niche preemption. Alternatively, we cannot rule out the hypothesis of both species being not spatially related. In contrast to Xcc8004, *S. rhizophila* is mainly located at the seed surface of radish, since no strains were recovered after seed surface sterilization (unpublished observations). Seed surface localization is in accordance with the epiphytic localization of *S. rhizophila* DSM 14405 on leaves and roots tissues of cotton, sweet pepper and tomato^52^. Owing to this predicted localization, it is tempting to postulate that *S. rhizophila* is a late colonist of the seed microbiome, being transmitted throug the external pathway.

Nowadays, sanitary quality of seeds is achieved through chemical methods and prophylaxis measures. Since reducing pesticide usage is an important objective for sustainable agriculture, the search for alternative seed treatments is essential. One of these alternative treatments consists in coating the seeds with microorganisms possessing biocontrol activities. However, biocontrol-based strategies require a better understanding of the colonization abilities of the microbial consortia within the seed habitat and its persistence within the spermosphere^21^. Our data showed that *S. rhizophila* and Xcc8004 have a high resource overlap and can compete *in vitro* for resources. An interesting approach to restrict seed transmission of Xcc8004 could be the introduction of *S. rhizophila* into the seed tissues with EndoSeed^TM^ technology^20^. Inoculation would encourage the co-existence of both taxa within the same ecological niche and maybe limit the size of Xcc8004 population within seed tissues. In addition to this augmentative biological control strategy, the limitation of Xcc8004 transmission could be potentially obtained by sowing radish seeds in plots containing a high-level of *S. rhizophila*. Indeed, the composition of the spermosphere-associated microbial assemblages is highly influenced by local horizontal transmission^53^. All things considered*, S. rhizophila* seems to be a promising candidate to reduce seed transmission of the pathogen Xcc8004, although further seed transmission experiments are needed to corroborate the fate of these taxa *in planta*. Collectively, these data may serve as a foundation for further research towards the design of novel biocontrol-based strategies on seed samples.

## Conclusions

Seed-transmission of phytopathogenic agents is significant for disease emergence and spread. In the present study, we have shown that the presence of Abra43 and Xcc8004 within radish seed samples did not modify the structure and function of the seed microbiome. The absence of community shift is explained in part by the low overlap in the use of resources between Xcc8004 and the main seed-borne bacterial populations. However, we have identified potential competitors of Xcc8004 belonging to *S. rhizophila* species. Understanding the pathways and determinants of seed transmission of Xcc8004 and *S. rhizophila* can result in the subsequent design of efficient management methods for restricting prevalence of phytopathogenic agents within seed samples through niche preemption^21^. More generally, the metagenomic sequences obtained in this work provide a starting point to investigate the genomic features involved in seed transmission through comparative genomics with bacterial isolates recovered from other plant habitats such as the rhizosphere or the phyllosphere^54^. Our approach of coupling community profiling approaches, bacterial confrontational assays and fluorescent *in situ* hybridization methods opens the way for further target research on seed inoculations with microorganisms possessing plant-beneficial properties.

## MATERIALS AND METHODS

### Seed collection, DNA extraction and shotgun metagenomic libraries preparation

The seed samples used in this work were collected in 2013 and 2014 during experiments thoroughly described elsewhere^24^. Briefly, radish seeds (*Raphanus sativus* var. Flamboyant5) were harvested at the FNAMS experimental station (47°28’12.42”N, 0°23’44.30”W, Brain-sur-l’Authion, France). These seed samples were collected from plants inoculated with *Alternaria brassicicola* strain Abra43; plants inoculated with *Xanthomonas campestris* pv. *campestris* strain 8004 and control plants without pathogen inoculation. To assess variation in seed community composition, every seed sample (A2013, A2014, C2013, C2014, X2013 and X2014) was divided into 3 subsamples (pseudo-replicates) of 10,000 seeds, resulting in a total of 18 samples (**Additional file 1**). In addition, a fourth subsample of 10,000 seeds was collected from the X2013 sample for PacBio sequencing.

DNA extraction was performed on seed samples according to the procedure described earlier^46^ (see **Text S1** in the Supplementary material for protocol details). DNA samples were sequenced at the Genome-Transcriptome core facilities (Genotoul GeT-PlaGe; France). Illumina sequencing libraries were prepared according to Illumina’s protocols using the Illumina TruSeq Nano DNA LT Library Prep Kit. DNA-sequencing experiments were performed on an Illumina HiSeq3000 using a paired-end read length of 2×150 bases with the Illumina HiSeq3000 Reagent Kit. Each library was sequenced on three lanes of HiSeq3000. PacBio sequencing was performed on a RSII system (details in **Text S1**). PacBio reads were used to improve the contigs length of the meta-assembly but were not used to assess differences in taxonomic/functional compositions. 0.25 nM of libraries were loaded per SMRTcell on the PacBio RSII System and sequencing was performed on 6 SMRTcells using the OneCellPerWell Protocol (OCPW) on C4P6 chemistry and 360 minutes movies.

### Metagenomic read processing

Sequencing of the 18 metagenomic seed samples resulted in a mean of 15.3 million paired-end reads per sample (SD ± 3.3 million). These paired-end reads were processed with cutadapt version 1.8^56^ to remove Illumina’s adapters, bases with Qscore < 25, reads possessing more than one ambiguity and paired-end reads with a length < 100 bases. Host-related reads were removed with Bowtie2 version 2.2.4^57^. Quality filtering of raw reads resulted in a mean of 12.5 million paired-end reads (SD ± 2.7 million) per metagenomics read set, which corresponds to approximately 2.5 Gb per metagenome (**Text S1** and **Additional file 1**).

### Taxonomic inference of metagenomics reads

The relative abundances of Xcc8004 and Abra43 within seed samples were estimated by mapping metagenomic reads to their respective genome sequences^23, 58^. Taxonomic profiling of each metagenome was estimated with a modified parallel version of Kraken^59^, named ParaKraken^33^, creating a count table of species and a relative abundance for each sample. Reads affiliated with the species *Xanthomonas campestris* and *Alternaria brassicicola* were removed in the count table for richness and diversity analyses (see **Text S1** for details).

Richness and diversity were calculated with the R package phyloseq 1.22.3^60^ on species count table. Differences in observed richness, estimated richness and diversity between samples were assessed through Kruskal-Wallis non parametric analysis of variance followed by Dunn’s post-hoc test. Differences in community membership and composition were estimated with the Jaccard and Bray-Curtis index. Principal coordinate analysis (PCoA) was used for ordination of Jaccard and Bray-Curtis distances. A permutational multivariate analysis of variance (PERMANOVA) was carried out to assess the impact of pathogen transmission on seed-associated microbial community profiles. To quantify the contribution of the seed transmission of phytopathogenic microorganisms on microbial community profiles, a canonical analysis of principal coordinates (CAP) was performed with the function capscale of the R package vegan 2.4.2, followed by PERMANOVA with the function adonis.

To investigate changes in the relative abundance between the different seed samples, rare species (those occurring in less than 75% of the samples) were first removed from the species count table. Fcros differential abundance analysis^31^ was then performed between each sample pair on all species and just the species with the highest abundance (>0.1% RA). A p-value cutoff of 0.05 and an f-score of 0.9 were used to determine differential abundance. The species scaled data was also run through DUO (https://github.com/climers/duo.git) with a threshold of >0.75, a program that identifies positive and negative relationships between species. Briefly, DUO first identifies abundance thresholds of 75^th^ and 25^th^ percentiles. The percentiles are then used to identify relationships between taxa by generating four categories (high-high, high-low, low-high, and low-low) based upon the CCC metric^32^.

### Functional profiling of each metagenome

An assembly of Illumina and PacBio metagenomic reads was performed with IDBA_UD^61^ and HGAP3^62^, resulting in a non-redundant assembly of circa 1 Gb containing 317,206 contigs (**Additional file 12**). A gene catalogue was created through gene prediction with Prokka 1.12^63^. Orthology inference of the 930,134 predicted CDSs was performed with DIAMOND^64^ against the EggNOG v4.5 database^65^. Approximately 73% of the predicted CDSs (682,374 CDSs) were affiliated with one orthologous group (OG).

To avoid an overrepresentation of orthologous groups (OGs) related to the inoculated phytopathogenic agents, reads that were mapped to Xcc8004 and Abra43 genomes were discarded. The remaining metagenomic reads were mapped with BowTie2 against all predicted CDSs and converted to bam files with Samtools v1.2.1^66^. Bam files were used to count the number of reads occurrence within each predicted CDS.

Similar to the species counts data, the OG counts were run through DUO with a threshold of >0.75 to identify positive (up-up and down-down) and negative (down-up and up-down) relationships between OGs to hypothesize which functional elements may interact. OG DUO networks were split into a network of positive interactions and negative interactions and then clustered using MCL^67^ to identify groups of highly interconnected OGs.

### Reconstruction of metagenome-assembled genomes

Initial binning was performed with Metabat v. 0.32.4^68^. Metagenome-assembled genomes completeness, contamination and strain heterogeneity were evaluated with CheckM v 1.0.4^69^. Taxonomic affiliation of each bin was performed by Average Nucleotide Identity based on BLAST (ANIb) with Jspecies^70^. Closest ANIb values were selected for reference genomes and used for Circle Packing representation with the D3.js JavaScript library representation (see details in **Text S1**).

### Reconstruction of genomic sequences of seed-associated bacterial isolates

Approximately 200 bacterial strains from the seed samples used for the metagenomics analysis were isolated on 1/10 strength Tryptic Soy Agar (**Text S1**). A total of 21 representative isolates of the main bacterial taxa were subjected to Illumina HiSeq4000 PE150 sequencing (**Table 1** and **Additional File 6**). These isolates were chosen by comparing their *gyrB* haplotypes with *gyrB* sequences previously obtained on radish seed samples^24^. All genome sequences were assembled with SOAPdenovo 2.04^71^ and VELVET 1.2.10^72^, annotated with prokka^63^ and EggNOG 4.5^65^.

### Characterization of resource overlap between seed-associated bacterial isolates

Nutritional resource consumption patterns of bacterial strains were assessed with Biolog GEN III MicroPlate^TM^ (Biolog Inc) (see details in **Text S1** in Supplementary data). One Biolog GEN III MicroPlateTM was used per isolate. The nutritional resources overlap was also predicted by comparing the numbers of OGs that are classified in the following functional categories: [E] amino acid transport and metabolism, [G] carbohydrate transport and metabolism, [I] lipid transport and metabolism and [P] inorganic ion transport and metabolism. Competition for resources and direct antagonisms between Xcc8004 and the selected bacterial strains were assessed on radish exudates and TSA 10% media (details in **Text S1**).

### Community profiling of germinating seeds and radish seedlings

Seed samples harvested in plot X2014 were used to determine the dynamics of bacterial population during germination and emergence of radish. DNA extraction and subsequent *gyrB* amplicon library preparation were performed on germinating seed (*n*=6) and seedling (*n*=8) samples according to the procedure described earlier^46^ (see details in **Text S1**). Libraries were sequenced with a MiSeq reagent kit v2 (500 cycles). Fastq files were processed with DADA2 1.6^73^ using the parameters described in the workflow for “Big Data: Paired-end”. The only modification made regarding this protocol was a change in the truncLen argument according to the quality of the sequencing run. Species abundance was assessed on germinating seed and seedling samples with the R package phyloseq^60^ (ASVs taxonomic affiliation details in **Text S1**).

### DOPE-FISH and CLSM microscopy of Xcc8004 infected seeds

Seeds were surface-sterilized, cut in half and fixed in a paraformaldehyde solution. DOPE-FISH was performed with probes from Eurofins (Austria) labeled at both the 5’ and 3’ end positions according to Glassner *et al.*^74^ using an EUBmix targeting all bacteria (EUB338, EUB338II, EUB338III) coupled with the fluorochrome Cy3^75, 76^, and a *Xanthomonas* spp. targeting probe (5’ -TCATTCAATCGCGCGAAGCCCG- 3’) coupled with Cy5^77^. A NONEUB probe^78^ coupled with Cy3 or Cy5 was also used independently as a negative control (further protocol details in **Text S1**). Samples were observed under a confocal microscope (Olympus Fluoview FV1000 with multiline laser FV5-LAMAR-2 HeNe(G) and laser FV10-LAHEG230-2) (see **Text S1** for image processing details). Pictures were cropped, and whole pictures were sharpened to better observe the image details. All experiments were repeated on eight seeds for each condition. Images presented in this publication are the average of colonization.

### Availability of data and material

The datasets supporting the conclusions of this article are available in the SRA database under the accession number PRJNA454584.

Whole Genome Shotgun projects have been deposited at GenBank under the accessions QFZV00000000-QGAM00000000.

All bacterial strains have been deposited at the CIRM-CFBP (https://www6.inra.fr/cirm_eng/CFBP-Plant-Associated-Bacteria; 10.15454/1.5103266699001077E12).

## Acknowledgements

The authors wish to thank Guillaume Chesneau for his help with bacterial competition assays, Laurent Noel for providing the strain *pTac*::GUS-GFP, the FNAMS for running field experiments, the ANAN platform (SFR QuaSav) for amplicon sequencing and BGI for sequencing the bacterial genomes. We would also like to thank BBRIC network for all the bioinformatic support.

## Authors’ contributions

GTC, BG, SC, DJ and MB designed the study and made substantial contributions to the analysis and interpretation of the results. GTC, BG, JP and MBr carried out bioinformatic analyses and participated actively in the interpretation of the results. SC performed FISH analyses. SR and AP performed all the microbiological analyses and Biolog profiles. AR and OB carried out the HiSeq and PacBio sequencing of the environmental DNA. GTC, BG and MB wrote the manuscript with input from the other authors. All authors read and approved the final manuscript.

## Conflict of Interest statement

The authors declare no conflict of interest.

## Funding

This research was supported by the grant awarded by the Region des Pays de la Loire (metaSEED, 2013 10080); RFI Objectif Végétal (DynaSeedBiome) and the AgreenSkills+ fellowship programme, which has received funding from the EU’s Seventh Framework Programme under grant agreement n° FP7-609398. This research used resources from the Oak Ridge Leadership Computing Facility at the Oak Ridge National Laboratory, which is supported by the Office of Science of the U.S. Department of Energy under Contract No. DE-AC05-00OR22725. This research was also supported by the Plant-Microbe Interfaces Scientific Focus Area in the Genomic Science Program, the Office of Biological and Environmental Research (BER) in the U.S. Department of Energy Office of Science, and by the Department of Energy, Laboratory Directed Research and Development funding (ProjectID 8321).

