## Supplementary data for "Differences in resources use lead to coexistence of seed-transmitted microbial populations"

**Supporting online material**

**Table of contents.**

**1. MATERIALS AND METHODS (Text S1)**

1.1 DNA extraction

1.2 Metagenomic read processing

1.3 Taxonomic inference of metagenomics reads

1.4 Functional profiling of each metagenome

1.5 Reconstruction of metagenome-assembled genomes

1.6 Reconstruction of genomic sequences of seed-associated bacterial isolates

1.7 Characterization of resource overlap between seed-associated bacterial isolates

1.8 Community profiling of germinating seeds and radish seedlings

1.9 DOPE-FISH and CLSM microscopy of Xcc8004 infected seeds

**2. REFERENCES**

**3. ADDITIONAL FILES**

**Text S1.**

### 1.MATERIALS AND METHODS

#### 1.1 DNA extraction and shotgun metagenomics libraries preparation

DNA extraction was performed on seed samples according to procedure described [1] . Briefly, seeds were transferred in 250 mL of PBS supplemented with 0.05% (v/v) of Tween® 20. Samples were incubated for 2 hours and 30 minutes at room temperature under constant agitation (140 rpm). One ml of the suspension was harvested for subsequent microbiological analyses. Suspensions were centrifuged (6000 x g, 10 min, 4°C) and pellets were used for DNA extraction. Total DNA was extracted with the PowerSoil DNA isolation kit (MoBio Laboratories) using the manufacturer's protocol. Due to the high requirement of DNA quantities for PacBio RSII library preparation, the fourth subsample of X2013 was extracted with NucleoSpin Food (Macherey-Nagel) according to the manufacturer's protocol.

PacBio sequencing was performed on a RSII system. The library was prepared according to PacBio's protocols with sample quantity and quality controls validated on Qubit, Nanodrop and fragment Analyser. The library was prepared from 2.6 µg of DNA using the SMRTBell template Prep Kit and no shearing was performed on the DNA. Using the BluePippin system, DNA fragments longer than 10 kb were selected.

#### 1.2 Metagenomic read processing

Plant-related reads were removed from the metagenomics reads with Bowtie2 version 2.2.4[2]. As the genomic sequence of the radish genotype (*Raphanus sativus* var. Flamboyant5) employed in this work is not available, we constructed an in-house *Brassicaceae* database composed of the genome sequences of *Arabidopsis thaliana* (GCA_000001735.1), *A. lyrata* (GCA_000004255.1), *Brassica oleracea* (GCA_000695525.1), *B. rapa* (GCA_000309985.1) and *R. sativus* (GCA_000801105.2).

#### 1.3 Taxonomic inference of metagenomics reads

Taxonomic profiling of each metagenome was estimated with a modified parallel version of Kraken[3], named ParaKraken. Each sample counts were scaled by the median of the total counts between each sample. Species level information was extracted and then counts for the same species were combined, creating a count table of species and relative abundance for each sample.

Fcros differential abundance analysis[4] was then performed between each sample pair on all species and just the species with the highest abundance (>0.1% RA). A pvalue cutoff of 0.05 and an f-score of 0.9 were used to determine differential abundance. To assess relationships between species, the non-rare species were further scaled within species, dividing each species-sample abundance by the maximum abundance for the given species. The species scaled data was also run through DUO (https://github.com/climers/duo.git) to both identify pathogen-microbiome interactions and interactions in the microbiome as a whole.

#### 1.4 Functional profiling of each metagenome

After mapping the metagenomics reads against all predicted CDSs, bam files were used to count the number of reads occurrence within each predicted CDS; The number of read per base was used as a measure of relative abundance of each orthologue group (OG). OG sample counts were then scaled by the median of the total counts between each sample. The OG scaled counts were further scaled dividing each OG sample count by the maximum count for the given OG.

#### 1.5 Reconstruction of metagenome-assembled genomes

Metagenomics reads of each seed sample were mapped to the meta-assembly using Bowtie2 and converted to bam file. Contigs composition and coverage were used for grouping contigs (> 5 kb) into metagenome-assembled genomes (MAGs) with Metabat v. 0.32.4 [5]. MAG completeness, contamination and strain heterogeneity were evaluated with CheckM v 1.0.4[6]. MAGs with a minimum of 50% completeness and less than 10% of contamination were selected for comparative genomics analyses. Gene prediction and functional annotations were performed with Prokka 1.2[7] and EggNOG 4.5[8]. Taxonomic affiliation of each bin was performed by similarity search (BLASTn[9]) of each contig against the nt database NCBI. The genome sequence corresponding to the best BLASTn hit of each contig was then retrieved to perform Average Nucleotide Identity based on BLAST (ANIb) with Jspecies[10]. Closest ANIb values were selected for references genomes and used for Circle Packing representation with the D3.js JavaScript library representation.

#### 1.6 Reconstruction of genomic sequences of seed-associated bacterial isolates

Approximately 200 bacterial strains from seed samples employed for metagenomics analysis were isolated on 1/10 strength Tryptic Soy Agar (17 g.l-1 tryptone, 3 g.l-1 soybean peptone, 2.5 g.l-1 glucose, 5 g.l-1 NaCl, 5 g.l-1 K2HPO4, and 15 g.l-1 agar) after 7 days of incubation at 18°C. These isolates were typed through sequencing of a portion of gyrB with the primer set gyrB_aF64 and gyrB_aR353 following the procedure described[1].

#### 1.7 Characterization of resource overlap between seed-associated bacterial isolates

Nutritional resource consumption patterns of bacterial strains were assessed with Biolog GEN III MicroPlateTM (Biolog Inc). Bacterial isolates were grown on TSA 10% for 24h. Bacterial colonies were resuspended in inoculating fluid A (IF A) at final turbidity of 95% and 100 µl of this suspension were inoculated in each well of the GEN III MicroPlateTM. After 72h of incubation, OD490nm was recorded. A well was considered as positive at OD490nm > 0.1 after blank subtraction. Positive wells were used to compare the nutritional resource utilization potential of Xcc8004 and the other seed-associate bacterial isolates. One Biolog GEN III MicroPlateTM was employed per isolate. We defined nutritional resources overlap as the number of shared nutritional resources between Xcc8004 and the tested bacterial strain divided by the total number of nutritional resources used by Xcc8004 and tested strain.

Competition for resources between Xcc8004 and the selected bacterial strains were assessed on radish exudates media. Seed exudates were collected by soaking 10 g of seeds (approximately 1,000 seeds) in 40 ml of sterile water for 2h 30 min at 4°C under constant agitation (140 rpm). The suspension was collected and sterilized with a 0.22 µm pore size filter (Whatman, Fischer Scientific, UK). Forty mL of the exudates suspension was mixed with 360 mL of water Agar 1.5% (w/v), resulting in radish exudates media. The strain Xcc8004 GUS, that expressed GUS and GFP via chromosomal integration of *pTac*::GUS-GFP[11], was mixed with selected bacterial strains at a 1:1 ratio ($OD_{600nm}$ = 0.1). Ten µl of the resulting bacterial suspensions were spotted on radish exudates media. Bacterial spots were removed 1, 2, 3, 4 and 5 days post-inoculation (dpi), diluted in PBS and plated on TSA 10% supplemented with rifampicin (50 $mg.l^{-1}$) and X-Gluc (50 $µg.l^{-1}$, ThermoFischer Scientific) for assessing the number of Xcc8004 colony-forming units (CFU). Differences in number of CFUs per treatment were assessed by one-way ANOVA with post-hoc Tukey's HSD test. Three independent biological replicates each consisting of three technical replicates were performed. Direct antagonisms between the bacterial strains and Xcc8004 were assessed by seeding Xcc8004 at a concentration of 1 x 107 CFU per ml-1 in TSA 10%. Three fresh colonies of each bacterial isolate were spotted on each agar plates. Antagonism was recorded when a discernible clearing zone was observed around the inoculated-colonies.

#### 1.8 Community profiling of germinating seeds and radish seedlings

Six subsamples of 100 seeds per sample were incubated in sterile plastic boxes and incubated at 20°C in obscurity for 24 and 72 hours after imbibition in sterile distilled water. Fifty germinating seeds (24h) and 25 seedlings (72h) were collected per sample for DNA extraction. DNA extraction and subsequent *gyrB* amplicon library preparation were performed on germinating seed (n=6) and seedling (n=8) samples according to procedure described earlier[1]. Briefly, amplification of *gyrB* was performed with the primer set gyrB_aF64

(5'-MGNCCNGSNATGTAYATHGG-3') and gyrB_aR353 (5'-ACNCCRTGNARDCCDCCNGA-3') using the following cycling reactions: 94°C (2 min), 35 cycles of amplification 94°C (30 s), 55°C (60 s) and 68°C (90 s) with a final extension step of 10 min at 68°C. All amplicons were purified with Agencourt AMPure XP system and quantified with QuantIT PicoGreen. A second round of amplification was performed with 5µl of purified amplicons and primers containing the Illumina adapters and indexes. PCR cycling conditions were: 94°C (2 min), followed by 12 cycles of amplification (94°C for 1 min, 55°C for 1min, 68°C for 1 min) and a final extension step at 68°C (10 min). Libraries were sequenced with a MiSeq reagent kit v2 (500 cycles).

Taxonomic affiliation of the amplicon sequence variants (ASVs) generated with DADA2[12] was performed with a naive Bayesian classifier on an in-house *gyrB* database. Other ASVs not classified at the phylum level were discarded from the datasets. The remaining ASV were rarefied at 50,000 reads per sample and agglomerated at the species level.

#### 1.9 DOPE-FISH and CLSM microscopy of Xcc8004 infected seeds

Seeds were surface sterilized by immersing seeds in 96% ethanol for 3 min, before being immersed in sodium hypochlorite (30g/L) for 2 min and rinsed thrice in sterile distilled water. They were then cut in half and fixed in a paraformaldehyde solution (4% w/v in PBS, pH 7.2) in Eppendorf tubes, before to be rinsed thrice with PBS. Samples were then treated with lysozyme (1 mg.mL-1 in PBS) during 10 min at 37C, rinsed thrice again and dehydrated in an ethanol series (50, 70, 99.9%; 15 min each step). DOPE-FISH was performed with probes from Eurofins (Austria) labeled at both the 5' and 3' end positions according to Glassner et al[13] using an EUBmix targeting all bacteria (EUB338, EUB338II, EUB338III) coupled with the fluorochrome Cy3[14,15], and a *Xanthomonas* spp. targeting probe (5' - TCATTCAATCGCGCGAAGCCCG- 3') coupled with Cy5[16] were used. A NONEUB probe[17] coupled with Cy3 or Cy5 was also used independently as a negative control. Hybridization was carried out by placing seed samples in Eppendorf tubes at 46°C for 2 h with 20 µL solution/per seed (containing 20 mM Tris–HCl pH 8.0, 0.01% w/v SDS, 0.9 M NaCl, 15% formamide, 10 ng µL-1 of each general probe and 5 ng µL-1 of specific probe). Post-hybridization was conducted at 48°C for 20 min with a pre-warmed post-FISH solution containing 20 mM Tris–HCl pH 8.0, 0.01% SDS, and NaCl at a concentration corresponding to the formamide concentration. Samples were then rinsed with distilled water before air drying for at least 1 day in the dark. The samples were then put on microscope slides and observed under a confocal microscope (Olympus Fluoview FV1000 with multiline laser FV5-LAMAR-2 HeNe(G) and laser FV10-LAHEG230-2). X, Y, Z pictures were taken at 405, 568, and 633 nm with 60X objectives. Affiliated colors on confocal equipment were green and red for Cy3 and Cy5, respectively, to separate well signals of Cy3 (normally yellow/greenish fluorescent) and Cy5 (red). Pictures were taken with analog integration or photon counting detection modes. Pictures were then observed using Imaris and Image J softwares[18]. Z Project Stacks were used to create the pictures (as described in [19]). Pictures were cropped, and whole pictures were sharpened to better observe the image details. All experiments were repeated on eight seeds for each condition. Images presented in this publication are the average of colonization.

### 2.REFERENCES


1. Barret, M. *et al.* Emergence shapes the structure of the seed microbiota. *Appl. Environ. Microbiol.* **81,** 1257–66 (2015).

2. Langmead, B. & Salzberg, S. L. Fast gapped-read alignment with Bowtie 2. *Nat. Methods* **9,** 357–359 (2012).

3. Wood, D. E. & Salzberg, S. L. Kraken: ultrafast metagenomic sequence classification using exact alignments. *Genome Biol.* **15,** R46 (2014).

4. Dembélé, D. & Kastner, P. Fold change rank ordering statistics: a new method for detecting differentially expressed genes. *BMC Bioinformatics* **15,** 14 (2014).

5. Kang, D. D., Froula, J., Egan, R. & Wang, Z. MetaBAT, an efficient tool for accurately reconstructing single genomes from complex microbial communities. *PeerJ* **3,** e1165 (2015).

6. Parks, D. H., Imelfort, M., Skennerton, C. T., Hugenholtz, P. & Tyson, G. W. CheckM: assessing the quality of microbial genomes recovered from isolates, single cells, and metagenomes. *Genome Res.* **25,** 1043–55 (2015).

7. Seemann, T. Prokka: rapid prokaryotic genome annotation. *Bioinformatics* **30,** 2068–2069 (2014).

8. Huerta-Cepas, J. *et al.* eggNOG 4.5: a hierarchical orthology framework with improved functional annotations for eukaryotic, prokaryotic and viral sequences. *Nucleic Acids Res.* **44,** D286–D293 (2016).

9. Altschul, S. F., Gish, W., Miller, W., Myers, E. W. & Lipman, D. J. Basic local alignment search tool. *J. Mol. Biol.* **215,** 403–410 (1990).

10. Richter, M., Rosselló-Móra, R., Oliver Glöckner, F. & Peplies, J. JSpeciesWS: a web server for prokaryotic species circumscription based on pairwise genome comparison.

*Bioinformatics* **32,** 929–931 (2016).

### 3.ADDITIONAL FILES

**Additional file1: Number of reads per metagenomics samples.** The number of raw paired-end (PE) reads, filtered PE reads, detected *Brassicaceae* PE reads, classified PE reads, mapped PE reads to Xcc8004 and Abra43 genomic sequences, respectively are indicated in the different columns. The percentage of filtered, classified and mapped reads in relation to the number of raw PE reads are also displayed.

**Additional file2: Taxonomic composition of the fungal fraction of the seed microbiome.** Estimation of the taxonomic composition of the fungal fraction of the radish seed microbiome was estimated with the ITS1 region of the fungal internal transcribed spacer (A) or k-mer based prediction performed on metagenomics samples (B).

**Additional file3.cys: Fcros differentially abundant species across seed samples.** Cytoscape session showing fcros analyses of microbial species detected on seed samples contaminated with Abra43 or Xcc8004 and control seed samples. *Pseudomonas* is the dominant genus that is differentially abundant across all conditions (files are in the share folder: https://www.dropbox.com/sh/98zlt2izpwf9hbq/AACxiR2ihMQF1Am8iOW1ayJVa?dl=0)

**Additional file4.cys: Positives DUO interactions between species during seed transmission of Xcc8004 and Abra43**. Cytoscape session showing the clusters of species with DUO scores >0.75. No strong DUO interactions were identified for Xcc8004 and Abra43, suggesting little overlap in niche space with other organisms within the microbiome conditions (files are in the share folder: https://www.dropbox.com/sh/98zlt2izpwf9hbq/AACxiR2ihMQF1Am8iOW1ayJVa?dl=0)

**Additional file5.cys: Potential pathogenesis-related subnetwork of OG DUO interactions.** DUO was run on the scaled OG counts and then clustered with MCL to identify highly interactive networks. Four networks are presented in the cytoscape session: the full positive network, full negative network, and the MCL networks for positive and negative network. One such network is a group of positively associated domains that have functions related to host-micobiome identification and immune activation, cell membrane modification for potential invasion, and host cell response to stress/invasion through production of preautophagosomal phagophore assembly site organisation. conditions (files are in the share folder: https://www.dropbox.com/sh/98zlt2izpwf9hbq/AACxiR2ihMQF1Am8iOW1ayJVa?dl=0)

**Additional file6: Phylogenetic relationships between seed-associated bacterial strains.** A distance tree (Neighbor joining) was calculated from partial *gyrB* sequence of bacterial isolates. Isolates in bold have been selected for genome sequencing. Grey bars represented the predicted occurrence of the isolates based in seed samples according to *gyrB* datasets.

**Additional file7: Relationship between genomic sequences**. Circle packing of ANIb values obtained from the different bacterial genomes can be visualized in the link: file:///home/gtorrescort/Documents/Radis_paper/FINALVERSION_Submisson_Mbio/submission/Additional_file7.html

**Additional file8: Average coverage of each genomic sequences.** Hierarchical clustering of seed samples based on the average coverage of the genomic sequences and metagenomics bins obtained in this work.

**Additional file9: GENIII profiles of selected bacterial strains.** Hierarchical clustering of bacterial strains based on their GENIII profiles. Metabolite reduced by a bacterial strain is indicated in red.

**Additional_file10.xlsx: Specific Xanthomonadales CDSs.** CDSs that are exclusively between Xcc8004 and the *S. rhizophila* strains sequenced in this work.

**Additional file11**: **Dynamics of the main bacterial species during germination and emergence of seed samples contaminated with Xcc 8004**. Amplification of *gyrB* was performed on DNA samples extracted on germinating seeds and seedlings. Samples were rarefied at 50,000 reads per sample. ASV detected with DADA2 were agglomerate at the species level. Colors represents the main bacterial families detected with *gyrB* during germination and emergence. Abundance is expressed in number of read per ASV. Every dot represents the relative abundance of ASVs per sample.

**Additional file12: Properties of metagenome assemblies**. The number of single reads, total number of base, number of contigs, maximum contig size, N50 and median contig size is indicated for each sample.

| Samples | Raw PE reads | Quality PE reads | | Brassicaceae PE reads | | Affiliated PE reads | | Xcc8004 PE reads | | Abra43 PE reads | |
|---|---|---|---|---|---|---|---|---|---|---|---|
| | counts | counts | % | counts | % | counts | % | counts | % | counts | % |
| A11_2013 | 19,970,262 | 16,679,514 | 83.52 | 50.387 | 0.25 | 15,391,943 | 92.56 | 36,370 | 0.22 | 104,870 | 0.63 |
| A11_2014 | 14,193,821 | 11,922,603 | 84 | 10.557 | 0.07 | 10,754,242 | 90.28 | 52,567 | 0.44 | 74,250 | 0.62 |
| A12_2013 | 20,700,090 | 17,079,882 | 82.51 | 97,850 | 0.52 | 15,342,081 | 90.34 | 34,374 | 0.2 | 239,552 | 1.4 |
| A12_2014 | 13,857,786 | 11,610,606 | 83.78 | 6.381 | 0.05 | 10,517,647 | 90.64 | 97,747 | 0.84 | 70,447 | 0.61 |
| A13_2013 | 19,336,327 | 15,952,078 | 82.5 | 89.224 | 0.46 | 14,398,487 | 90.76 | 50,234 | 0.32 | 192,376 | 1.21 |
| A13_2014 | 16,430,779 | 13,183,688 | 80.24 | 9.778 | 0.06 | 11,825,297 | 89.76 | 103,316 | 0.78 | 65,363 | 0.5 |
| C11_2013 | 10,310,051 | 8,190,680 | 79.44 | 135.923 | 1.32 | 6,486,388 | 80.52 | 127,079 | 1.58 | 12,955 | 0.16 |
| C11_2014 | 11,821,508 | 9,959,720 | 84.25 | 10.158 | 0.09 | 8,934,166 | 89.8 | 121,173 | 1.22 | 2,043 | 0.02 |
| C12_2013 | 14,297,360 | 11,622,310 | 81.29 | 163.561 | 1.14 | 9,673,822 | 84.42 | 139,799 | 1.22 | 13,522 | 0.12 |
| C12_2014 | 9,346,108 | 7,961,711 | 85.19 | 14.221 | 0.15 | 7,062,921 | 88.86 | 64,882 | 0.82 | 2,352 | 0.03 |
| C13_2013 | 13,242,052 | 10,778,736 | 81.4 | 174.115 | 1.31 | 8,742,507 | 82.44 | 76,122 | 0.72 | 15,853 | 0.15 |
| C13_2014 | 18,307,132 | 15,042,908 | 82.17 | 15.859 | 0.09 | 13,427,229 | 89.36 | 207,332 | 1.38 | 3,205 | 0.02 |
| X11_2013 | 17,966,998 | 14,537,543 | 80.91 | 221.003 | 1.23 | 12,248,293 | 85.56 | 754,427 | 5.27 | 16,299 | 0.11 |
| X11_2014 | 12,110,331 | 9,527,329 | 78.67 | 2.908 | 0.02 | 9,158,751 | 96.16 | 2,692,480 | 28.27 | 529 | 0.01 |
| X12_2013 | 17,797,945 | 13,882,096 | 78 | 134.949 | 0.76 | 12,361,651 | 89.92 | 1,213,018 | 8.82 | 8,201 | 0.06 |
| X12_2014 | 14,855,272 | 11,889,042 | 80.03 | 2.036 | 0.01 | 11,345,650 | 95.44 | 3,863,936 | 32.5 | 495 | 0.01 |
| X13_2013 | 16,667,027 | 13,675,232 | 82.05 | 85.963 | 0.52 | 12,319,799 | 90.66 | 504,548 | 3.71 | 12,015 | 0.09 |
| X13_2014 | 14,689,997 | 11,980,662 | 81.56 | 2.764 | 0.02 | 11,520,796 | 96.18 | 3,762,193 | 31.41 | 377 | 0.01 |

**Additional file1: Number of reads per metagenomics samples.**

**Additional file2: Taxonomic composition of the fungal fraction of the seed microbiome.**

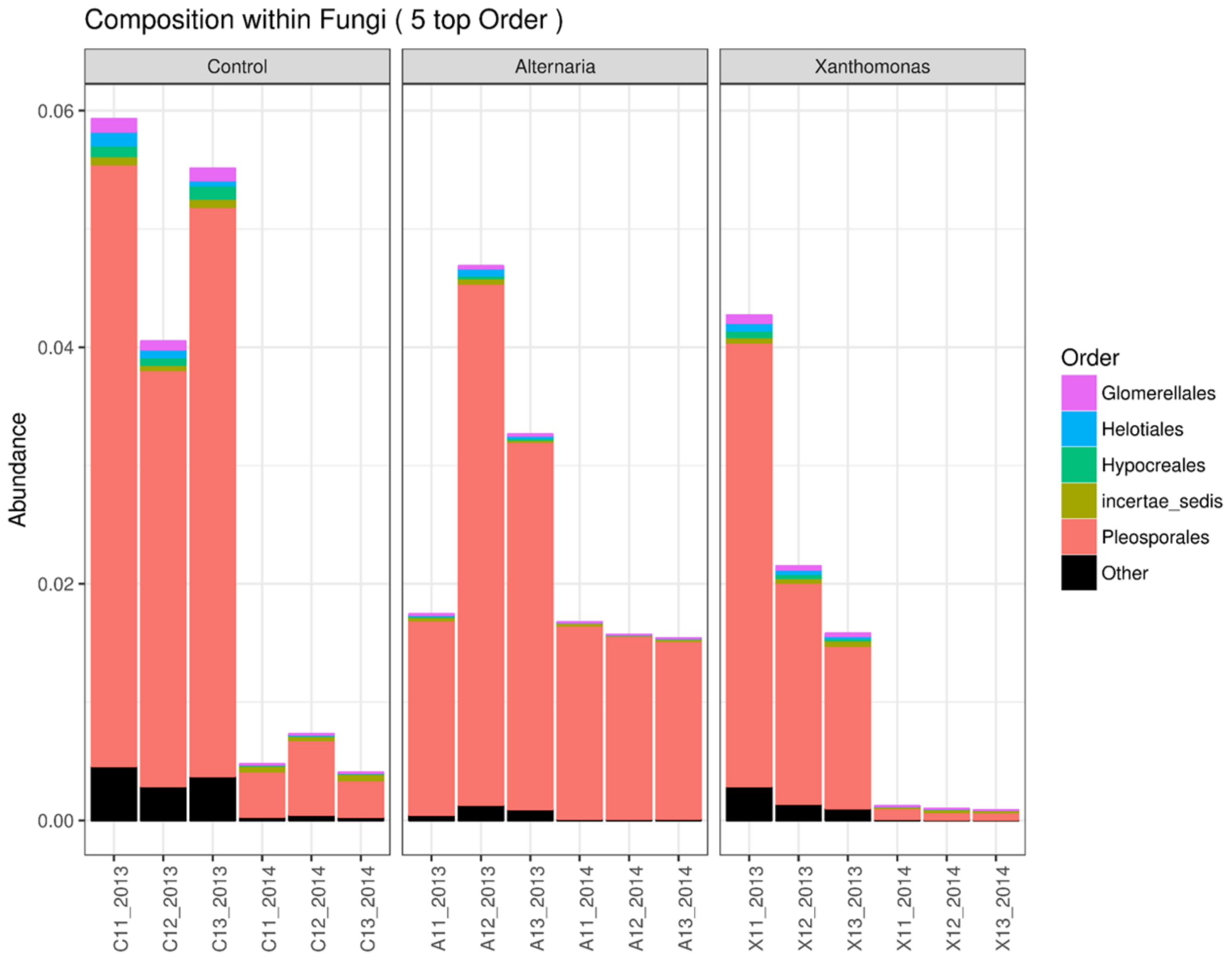

**Additional file6: Phylogenetic relationships between seed-associated bacterial strains.**

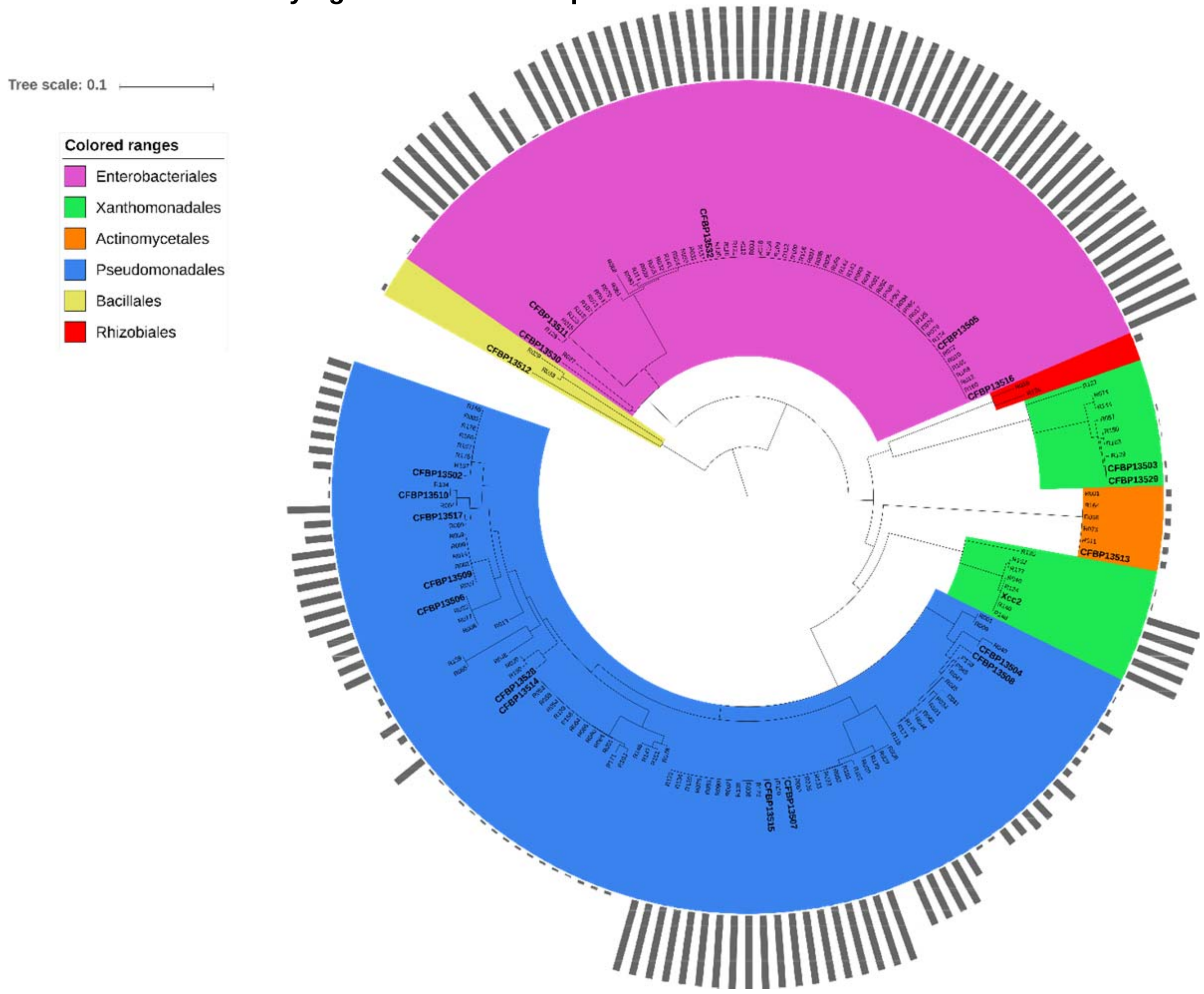

**Additional file8: Average coverage of each genomic sequences.**

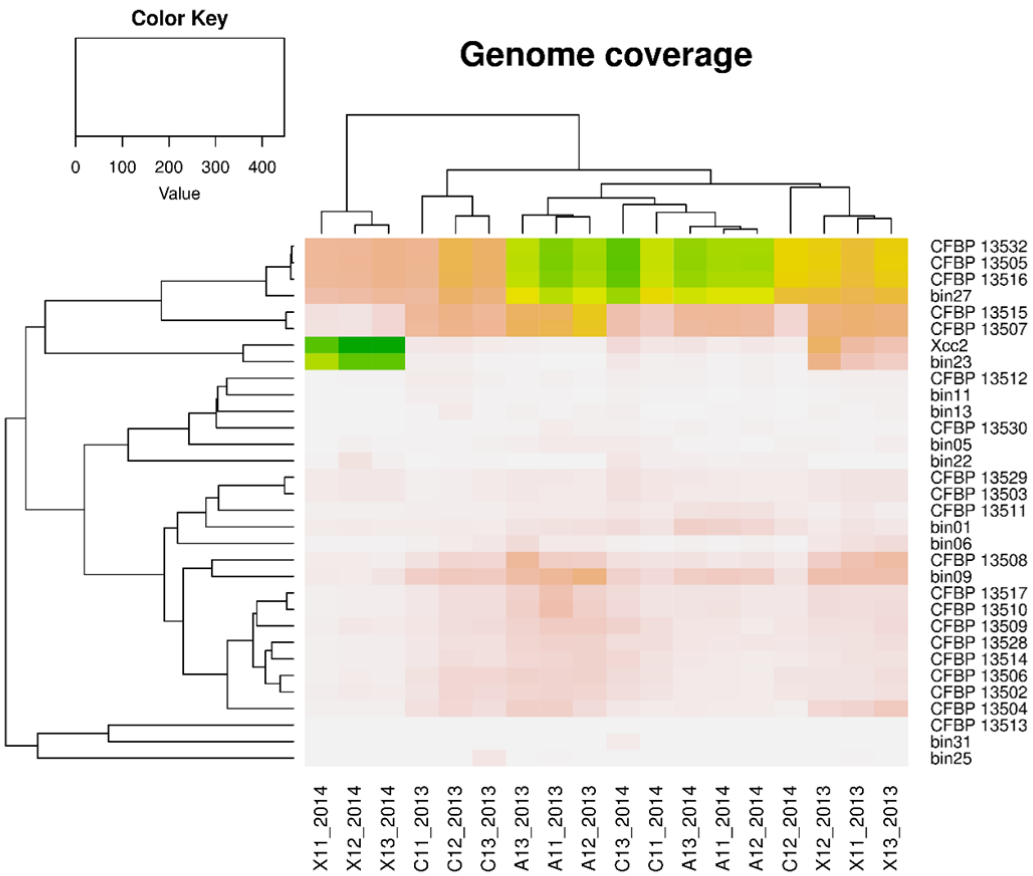

**Additional file9: GENIII profiles of selected bacterial strains.**

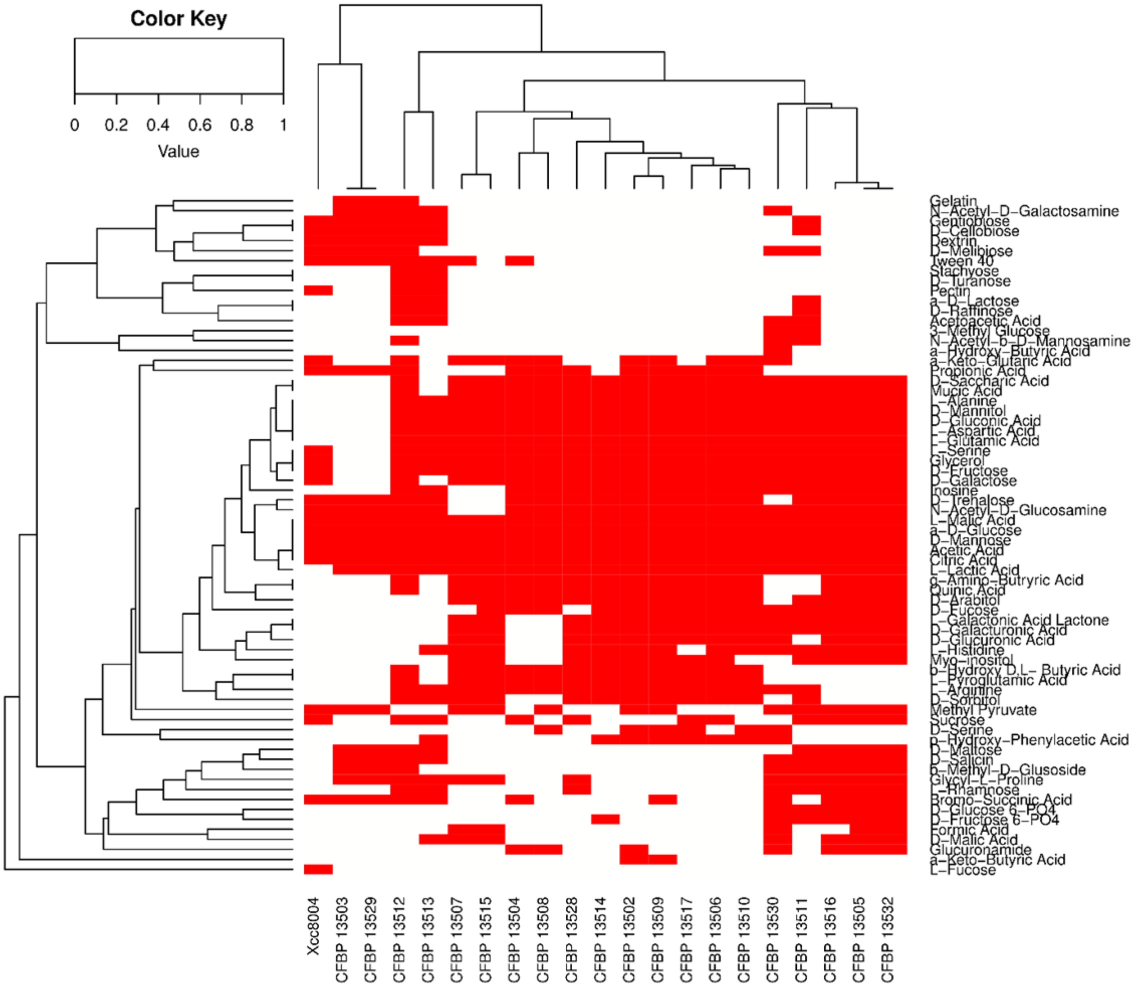

**Additional file11**: **Dynamics of the main bacterial species during germination and emergence of seed samples contaminated with Xcc 8004**.

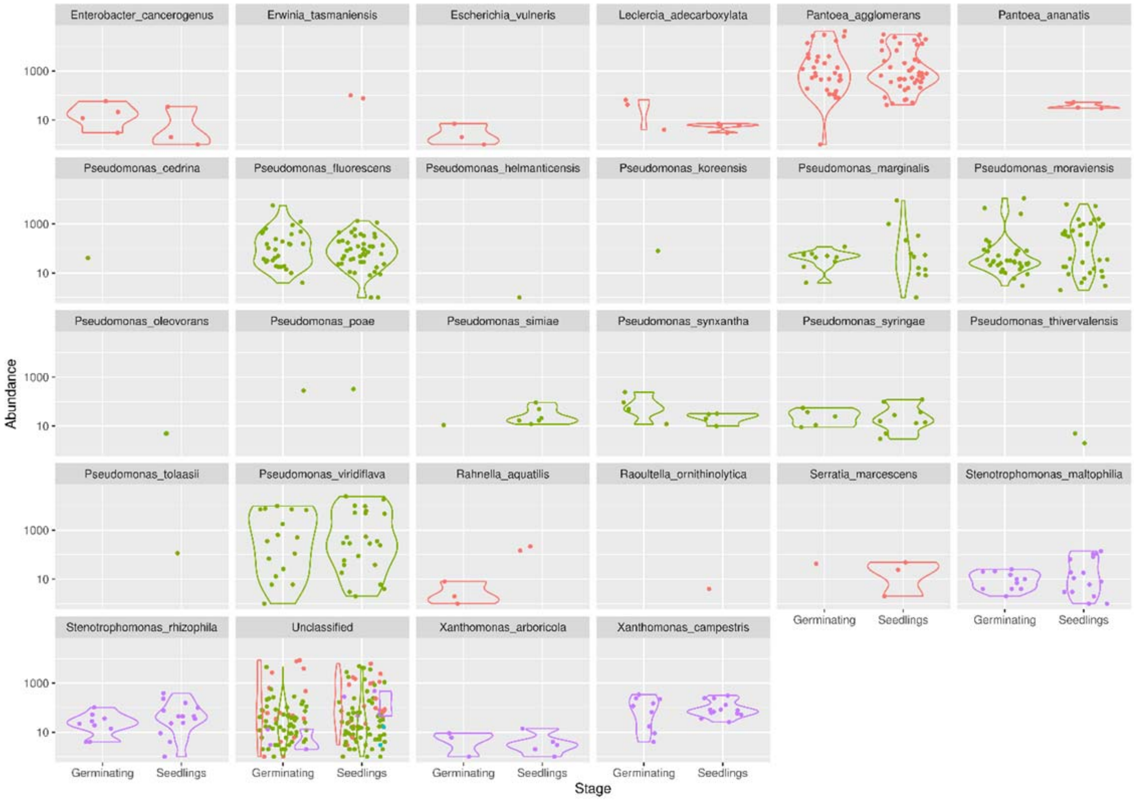

| Samples | Total reads | Base | Num | Max | N50 | Median |
|---|---|---|---|---|---|---|
| A11.2013 | 33,258,254 | 74,102,387 | 27.337 | 84.429 | 3.115 | 1.879 |
| A11.2014 | 23,824,092 | 38,667,011 | 14.899 | 66.512 | 2.842 | 1.907 |
| A12.2013 | 33,964,064 | 79,631,123 | 30.986 | 92.577 | 2.831 | 1.846 |
| A12.2014 | 23,208,450 | 40,339,142 | 15.082 | 43.46 | 3.021 | 1.933 |
| A13.2013 | 31,725,708 | 71,981,192 | 23,420 | 133.896 | 3.863 | 1.947 |
| A13.2014 | 26,347,820 | 50,615,061 | 17.877 | 89.372 | 3.273 | 2.059 |
| C11.2013 | 16,109,514 | 39,218,266 | 12.804 | 119.973 | 3.939 | 1.783 |
| C11.2014 | 19,899,124 | 49,241,376 | 15.837 | 87.954 | 3.786 | 2.049 |
| C12.2013 | 22,917,498 | 68,940,449 | 22.905 | 125.845 | 3.616 | 1.912 |
| C12.2014 | 15,894,980 | 34,101,903 | 12.298 | 50.834 | 2.775 | 1.926 |
| C13.2013 | 21,209,242 | 65.286677 | 20.203 | 254.355 | 4.126 | 1.869 |
| C13.2014 | 30,054,098 | 80,146,963 | 26.918 | 64.626 | 3.534 | 2.109 |
| X11.2013 | 28,633,080 | 56,340,707 | 20,460 | 167.217 | 3.125 | 1.782 |
| X11.2014 | 19,048,842 | 23,513,580 | 5.419 | 222.503 | 7.784 | 2.024 |
| X12.2013 | 27,494,294 | 52,144,072 | 17.089 | 248.629 | 3.667 | 1.857 |
| X12.2014 | 23,774,012 | 30,160,879 | 4.848 | 222.503 | 11.316 | 2.857 |
| X13.2013 | 27,178,538 | 57,843,992 | 17.668 | 222.505 | 4.286 | 1.925 |
| X13.2014 | 23,955,796 | 25,052,063 | 6.665 | 222.503 | 5.405 | 1.909 |
| all | 448,497,406 | 937,326,843 | 312,715 | 254.355 | 3.578 | 1.924 |

**Additional file12: Properties of metagenome assemblies**.
